# Signals of variation in human mutation rate at multiple levels of sequence context

**DOI:** 10.1101/385096

**Authors:** Rachael C. Aikens, Kelsey E. Johnson, Benjamin F. Voight

## Abstract

Our understanding of mutation rate helps us build evolutionary models and make sense of genetic variation. Recent work indicates that the frequencies of specific mutation types have been elevated in Europe, and that many more, subtler signatures of global polymorphism variation may yet remain unidentified. Here, we present an analysis of the 1,000 Genomes Project (phase 3), suggesting additional putative signatures of mutation rate variation across populations and the extent to which they are shaped by local sequence context. First, we compiled a list of the most significantly variable polymorphism types in a cross-continental statistical test. Clustering polymorphisms together, we observed four sets of substitution types that showed similar trends of relative mutation rate across populations, and describe the patterns of these mutational clusters among continental groups. For the majority of these signatures, we found that a single flanking base pair of sequence context was sufficient to determine the majority of enrichment or depletion of a mutation type. However, local genetic context up to 2-3 base pairs away contributes additional variability, and helps to interpret a previously noted enrichment of certain polymorphism types in some East Asian groups. Building our understanding of mutation rate in this way can help us to construct more accurate evolutionary models and better understand the mechanisms that underlie genetic change.

## INTRODUCTION

The process of mutation is a formative force in molecular evolution because it generates the genetic variation that can be acted upon by natural selection. Quantitative and qualitative insights regarding the mutation rate in human populations can facilitate the construction of increasingly informative models of human evolutionary history, targets of natural selection, and perhaps even genetic or environmental mechanisms that confer genomic stability and drive genetic change. While the rates of DNA mutation and repair have been known to differ widely between certain individuals^1^, within and between chromosomes^2^, and down to specific local sequences^3^, the biological mechanisms underlying mutation rate variability across the genome are not completely understood. Recent work has suggested that the mutation rate in humans may have been in flux over recent evolutionary history^4–6^. A key observation supporting this hypothesis is that the relative proportions of certain types of polymorphisms vary across populations. Most notably, studies cite the strong enrichment of <u>C</u>→T substitutions at certain trinucleotide contexts in Europe and South Asia^4–6^. These reports have also documented additional polymorphism types that appear to vary in frequency across populations^5,6^.

However, there are two lines of inquiry that have not yet been fully explored in these data. First, because clusters of polymorphisms with similar global profiles of enrichment might be driven by a shared mechanism, is worthwhile to ask not only which polymorphism types vary across the globe but also how variable polymorphism types group together as putative “profiles” of mutation rate variations. These profiles can then be analyzed together, for example by searching for ‘pulses’ of increased substitution volume across the site frequency spectrum which may indicate the timing of mutation rate changes in the distant past^6^. Developing a better understanding of these signatures of polymorphism variation may help to link such changes to a putative genetic or environmental cause. Second, models of local genetic sequence context beyond a single flanking nucleotide have not yet been applied to the study of variability in mutation rates across populations. Our previous work has demonstrated that windows of flanking genetic sequence broader than the trinucleotide context may uncover additional detail in mutation rate variability across the genome^3,7^. The consideration of greater numbers of upstream and downstream base pairs of genetic context could help to interpret the mechanism by which mutation rate varies across populations. For example, strong effects stemming from more remote sequence context may drive signals of mutation rate variation observed in a lower order (*i.e.*, trinucleotide) context, indicating that the underlying mechanism may rely on the broader local nucleotide configuration. As such, models that consider broader windows of local context may highlight subtle variation in polymorphism that might not have otherwise been detected.

For these reasons, we sought to expand upon previous studies by identifying polymorphisms at multiple context levels and quantifying how they vary across populations. To this end, we have applied a combination of sequence context frameworks to analyze the current release of the 1,000 Genomes project, spanning >2,000 subjects across four continents. Herein, we aimed to catalog population-level heterogeneity in polymorphism across the spectrum of sequence contexts, and cluster variable polymorphism classes that we observe into common mutational signatures which can be analyzed together across populations.

## RESULTS

To quantify differences in the frequencies of mutation types across populations, we assembled sets of genetic variants specific to Africans, Europeans, South Asians, and East Asians (504, 503, 489, and 504 individuals, respectively, excluding recently admixed American populations) from the 1000 Genomes Project (phase 3)^8^. As genetic variants in the coding genome are likely to be under purifying selection, we focused on variants observed in the non-coding genome to minimize the influence of selective pressures (**Methods**). Our final sets consisted of 7 million, 1.2 million, 1.96 million and 1.99 million variants private to African, European, East Asian and South Asian populations, respectively.

### Identifying novel 3-mer substitution classes that vary across continents

We first compiled a list of polymorphisms in trinucleotide (*i.e.,* ‘3-mer’) contexts that appear heterogeneous in their representation across the globe. To this end, we extended a previously described^4,6^ approach for comparing counts of polymorphisms between pairs of populations (**Methods**). Rather than making all possible pairwise comparisons between ancestral groups for a given substitution class, we perform a single test of homogeneity in private polymorphism for that substitution class across Africa, Europe, East Asia, and South Asia. In addition to reducing the required number of hypothesis tests compared to previous methods (important later for analyses with broader windows of sequence context), this statistical framework allowed us to rank order polymorphism types by the significance of their variation across all ancestral groups. After replicating previous results^4,6^ as a technical control (**Supplementary Fig. 1**), we applied our test to each 3-mer polymorphism type (96 total), applying a modified p-value correction (P_ordered_) as previously described^6^, in addition to Bonferroni correction for multiple hypothesis testing (nominal significance threshold P_ordered_ < 5 x 10^-4^).

As expected, the most compelling group of substitution classes was composed of <u>C</u>→T polymorphism types previously reported to be enriched in Europe and South Asia (**Table 1**). All four types that have been previously noted as part of this signal: T<u>C</u>C→T, A<u>C</u>C→T, T<u>C</u>T→T, and C<u>C</u>C→T were among the 6 most variable polymorphisms (all P_ordered_ < 1 x 10^-68^). We further observed that all four CpG transition mutations were variable between populations (P_ordered_ < 1 x 10^-31^, **Table 1, Supplementary Fig. 2**). Previous work has suggested that proportions of CpG substitutions are weakly variable between populations^5^. In agreement with this, we noted that all CpGs appear to have a shared profile of enrichment in South and East Asia, but that these ancestral variations in CpG substitution are small relative to the large overall abundance of C/T polymorphism at CpG sites. Importantly, Mathieson and Reich^5^ have cautioned that an apparent CpG enrichment may be a signature of recurrent mutation in populations which have experienced recent exponential growth^5^. However, if this were the case, we would expect to see a strong excess of CpG polymorphisms at doubletons (i.e., allele count = 2), which was not immediately apparent (****Figure 5C****).

**Table 1:** A cross-continental test for polymorphism variability at the 3-mer level. 15 polymorphism classes with 3-mer sequence contexts, shown here, were highly significant (P_ordered_ < 2 x 10^-37^) according to a chi-squared test for heterogeneity across non-admixed continental groups. Boldface numbers indicate a significant difference in polymorphism proportion compared with Africa (P < 1 x 10^-5^) in a pairwise chi squared test. P_ordered_ was calculated as previously described^6^ (**Methods**). *To facilitate comparison, approximate private mutation rates (per generation per site) for each continent were inferred by normalizing estimate substitution probabilities using all private mutations to the *de novo* mutation rate estimated from Kong et al^9^, and then subsequently normalized relative to inferred rate in Africa.

| Notes | 3-mer | Relative rate<br>in Africa* | Relative rate<br>in Europe* | Relative rate<br>in South Asia* | Relative rate<br>in East Asia* | P<sub>ordered</sub> |
| --- | --- | --- | --- | --- | --- | --- |
| Previously<br>reported C→T<br>elevation in<br>Europe<sup>4-6</sup> | T$\underline{C}$C→T | 1 | **1.56** | **1.20** | 1.00 | ≈ 0 |
| | A$\underline{C}$C→T | 1 | **1.20** | **1.07** | **0.93** | 3×10<sup>-308</sup> |
| | T$\underline{C}$T→T | 1 | **1.17** | **1.06** | 0.98 | 3×10<sup>-196</sup> |
| | C$\underline{C}$C→T | 1 | **1.06** | **1.03** | **0.95** | 2×10<sup>-69</sup> |
| CpG transition | T$\underline{C}$G→T | 1 | 1.02 | **1.06** | **1.04** | 2×10<sup>-49</sup> |
| | A$\underline{C}$G→T | 1 | 1.01 | **1.04** | **1.03** | 2×10<sup>-48</sup> |
| | G$\underline{C}$G→T | 1 | 1.01 | **1.04** | **1.04** | 4×10<sup>-45</sup> |
| Not previously<br>highlighted | G$\underline{A}$T→T | 1 | **1.06** | **1.13** | **1.21** | 5×10<sup>-111</sup> |
| | A$\underline{C}$C→A | 1 | **1.04** | **1.10** | **1.15** | 1×10<sup>-98</sup> |
| | A$\underline{C}$A→T | 1 | **0.95** | 0.97 | **0.93** | 3×10<sup>-60</sup> |
| | T$\underline{C}$A→T | 1 | **1.06** | **1.03** | 0.97 | 9×10<sup>-52</sup> |
| | A$\underline{C}$T→T | 1 | 1.02 | 1.00 | **0.94** | 3×10<sup>-51</sup> |
| | G$\underline{C}$T→T | 1 | 1.02 | **1.04** | **1.06** | 2×10<sup>-40</sup> |
| | G$\underline{A}$C→T | 1 | 1.02 | **1.09** | **1.20** | 6×10<sup>-40</sup> |
| | G$\underline{C}$C→T | 1 | **1.06** | 1.03 | 1.02 | 1×10<sup>-37</sup> |

We next examined the remaining variable polymorphisms for novel signatures of mutation rate variation. Surprisingly, we found that 63 of the 96 possible 3-mer types exceeded our Bonferroni correction for multiple tests across ancestral continental groups. Therefore, we opted first to consider only the top 15 most heterogeneous polymorphism types (corresponding to a P_ordered_ < 2 x 10^-37^, **Table 1**). To facilitate comparison of the relative enrichment between continental groups, we used all private polymorphisms to infer a mutation rate for each 3-mer (per generation per site) normalized to the average estimated *de novo* mutation rate from Kong et. al^9^ (**Methods**).

In addition to the <u>C</u>→T polymorphisms mentioned above, we observed eight additional polymorphism classes in our top ranked set that have not yet been specifically noted in previous studies (**Table 1**). One polymorphism, T<u>C</u>A→T, showed enrichment in Europe and South Asia, a profile similar to the previously noted <u>C</u>→T polymorphism types elevated in Europeans (**Table 1**). The pattern of enrichment for A<u>C</u>T→T was similar to this, with greatest representation in Europe and South Asia, but with a higher proportion in Africa compared to other substitution classes from this group. A final polymorphism, G<u>C</u>C→T, was likewise enriched in Europe and South Asia, but also showed an elevation in East Asia. Interestingly, G<u>A</u>T→T, A<u>C</u>C→A, G<u>A</u>C→T, and G<u>C</u>T→T polymorphisms displayed a similar profile of heterogeneity distinct from previously reported signatures of variation: the highest rates in East and South Asia, with intermediate levels in Europe, relative to Africa. This suggests that these three substitution classes may represent a group of polymorphism types enriched in Asia. A final substitution class, A<u>C</u>A→T, was most elevated in Africans, with lower rates in East Asians and Europeans, perhaps suggestive of an Africa-enriched signature.

As an additional validation, we estimated the mutation rate separately on each chromosome and found that these patterns of enrichment hold relatively consistently across the genome for each of these substitution classes (**Supplementary Fig. 3**). Taken together, these results support the proposition that there may be several previously unreported signatures of variation in mutation rate observed at the 3-mer scale, beyond the previously reported signal of European <u>C</u>→T enrichment.

### Hierarchical clustering of 3-mer mutational signatures

We next sought to identify sets of substitution classes that share similar patterns of enrichment or depletion across the globe, which we hypothesize might be influenced by a common underlying mechanism. To this end, we performed hierarchical clustering of 3-mer polymorphism types based upon their relative inferred mutation rates in each of the twenty 1,000 Genomes Project subpopulations comprising the non-admixed continental groups from our initial analysis.

We highlight five distinctive patterns of substitution rates - denoted here as “profiles” - that emerged from the clusters of 3-mer substitution classes (**Figure 1A**). Profile #1 corresponds to an enrichment of <u>C</u>→T substitutions in Europe, and includes all four 3-mers previously reported to comprise this signal^5,6^ (**Figure 1B**). The remaining three polymorphisms in this group, T<u>C</u>A→T, A<u>C</u>T→T, and G<u>C</u>C→T are noted in the previous section (**Table 1**), and represent variable polymorphisms not previously reported to be part of the Europe-enriched <u>C</u>→T signature.

**Figure 1:**
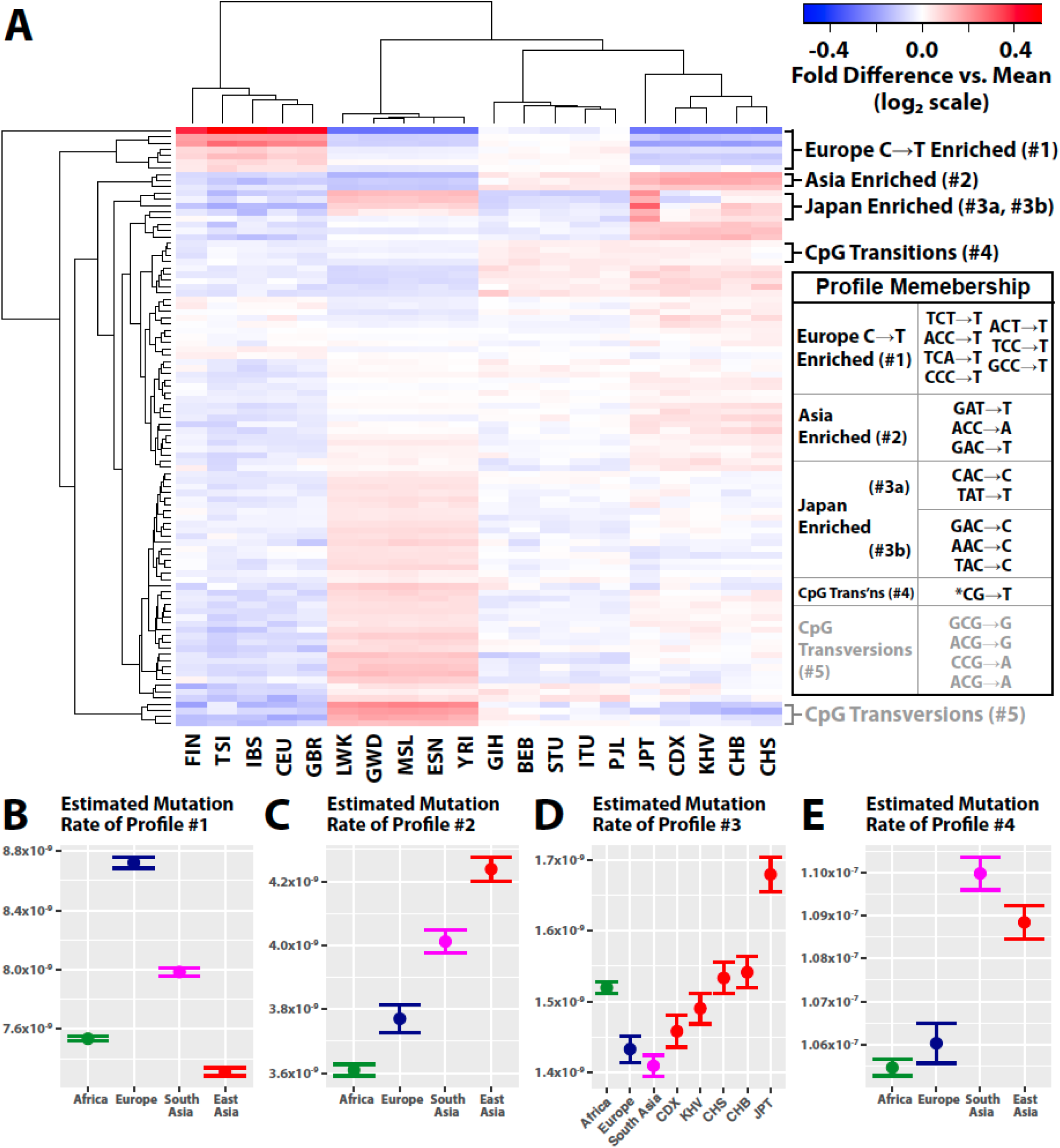
Profiles of mutation rate by trinucleotide context highlight variability across populations. **(A)** Heat map of all 3-mer polymorphisms, clustered based on their relative rates in each of twenty non-admixed 1,000 Genomes Project populations. Clusters of interest are labeled, and their membership is detailed in the table to the right. Polymorphisms are clustered and colored based on fold elevation over the mean mutation rate for each mutation type. All units are log base 2 transformed, with red color corresponding to enrichment and blue to depletion. Population codes along the bottom correspond to the three-letter codes assigned by 1,000 genomes **(B, C, E)** Approximate 95% confidence interval estimates of inferred mutation rate across continental groups for profiles 1-3. **(D)** Inferred mutation rates for profile 3 shown across Europe, Africa, South Asia, and five East Asian subpopulations: Chinese Dai in Xishuangbanna (CDX), Vietnamese (KHV), Han Chinese from Beijing and Southern China (CHB and CHS), and Japanese in Tokyo (JPT).

The next profile (#2) consists of G<u>A</u>T→T, A<u>C</u>C→A, and G<u>A</u>C→T, which are elevated in East and South Asia (**Figure 1C**). All three of these were noted above for their cross-continental heterogeneity (P_ordered_ < 7 x 10^-40^, **Table 1**). Next, we observed two clusters (Profiles #3a and #3b) that appear enriched in Japan and other populations in East Asia, relative to other continental groups. This unit is comprised of the *<u>A</u>C→C polymorphisms, as well as A<u>T</u>A→A, corresponding with a previous report^6^ which documented that *AC→C, and TAT→T mutation types separate East Asians in a principal component analysis. Together, profiles #2 and #3 may represent two distinct signatures of enrichment for certain mutation types in Asia.

The remaining two profiles both involve substitutions within CpG contexts. Profile #4, corresponds to the CpG transitions, which cluster together even after the data are normalized to show only relative mutation rates (**Figure 1D**). The final profile (#5) is comprised of CpG transversions that appeared to be elevated in Africa. However, Harris and Pritchard^6^ report that the proportions of these polymorphism types do not appear to agree between the Simons Diversity Genome Project and the 1,000 Genomes Project. This suggests that this profile may be the result of an experimental artifact, rather than a true divergence in mutation rate. In sum, the clusters identified here highlight sets of polymorphisms whose relative proportions are similar across populations encompassed by the 1,000 Genomes Project.

### Broader sequence contexts around 3-mer signatures

Our previous work has indicated that flanking nucleotides within broader windows of sequence context around a genetic locus strongly correlate with the probability of substitution^3,7^. Thus, we next took each 3-mer type identified as heterogeneous in the previous section and measured the frequency of private substitutions in a broader window of local sequence context that considered three flanking nucleotides (*i.e.*, a heptanucleotide, or ‘7-mer’ context window). This subdivided each 3-mer substitution into 256 distinct 7-mer classes, allowing us to ask whether the population-specific heterogeneity was common to all 7-mer expansions, or confined to a specific subset of those contexts. To this end, we plotted the relative inferred mutation rates of those polymorphisms in pairs of populations. If there were no signal of mutation rate difference between populations, we would expect all 7-mer expansions to be distributed along the y = x diagonal (case I, *e.g*., **Figure 2A**). If the most important local features driving a mutational signal lay within a single nucleotide base of the substitution, then we would expect all 7-mers to lie together off the diagonal (case II, *e.g*., **Figure 2B**). Alternatively, if a 3-mer signal were actually driven by a handful of highly variable 7-mer substitution types, only a handful of exceptional 7-mer types would lie far from the y = x line (case III).

**Figure 2:**
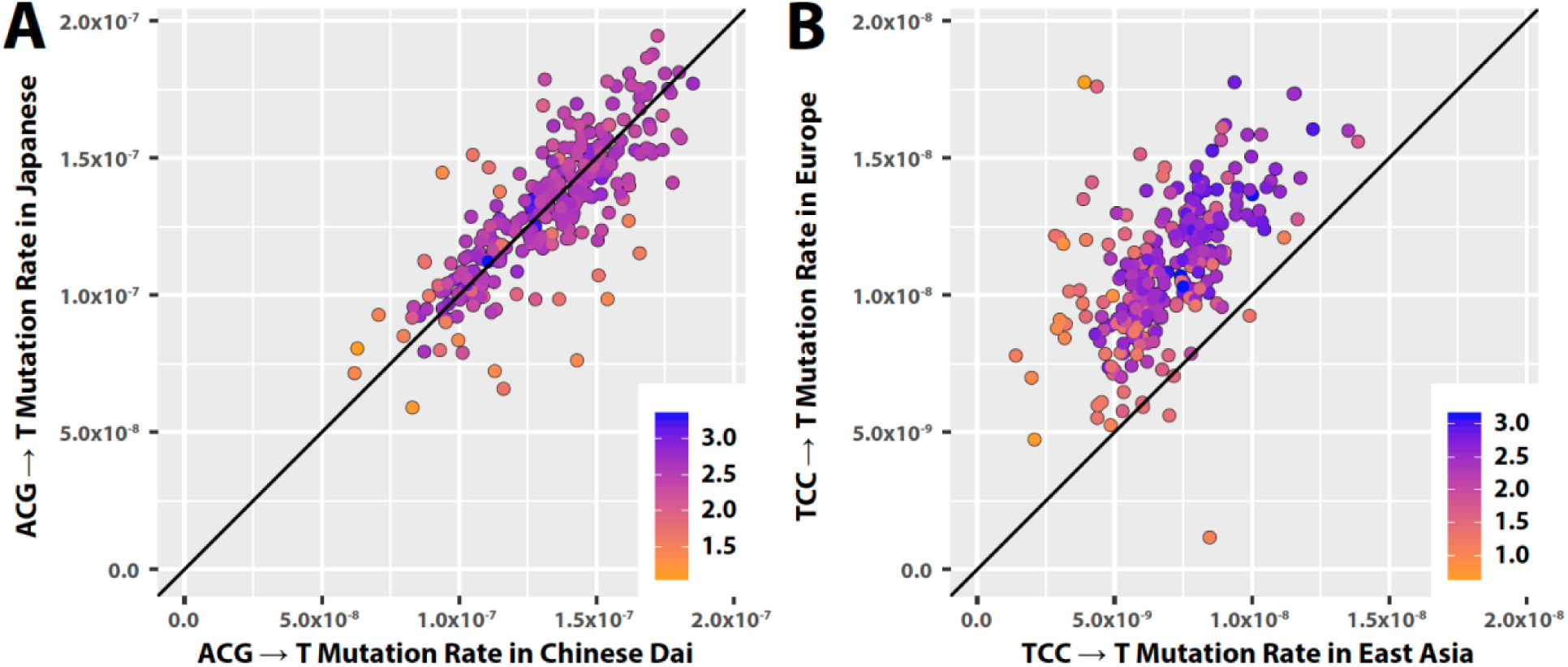
Mutational signatures driven at the 3-mer context level. Each point represents a 7-mer expansion of the 3-mer subtype shown, plotted based on its estimated mutation rate in each of the two populations displayed. Colors indicate the log (base 10) of the number of substitutions observed for that 7-mer class. **(A)** When a 3-mer substitution type occurs at equal rates in two related populations, most of the 256 7-mer expansions of this 3-mer appear along the diagonal y=x line. **(B)** For T<u>C</u>C→T and the other <u>C</u>↑T polymorphism types elevated in Europe, the bulk of the 7-mer expansions lie above the y=x diagonal, indicating that there has been a substantial difference in mutation rate between Europe and East Asia, and this difference is driven by effects at the 3-mer, rather than the 7-mer level.

We found that nearly all of the 3-mers comprising profiles #1, #2, and #4 matched case II (**Figure 2B**, **Supplementary Fig. 4-6**). Thus, the global variation in European <u>C</u>→T elevation, the CpG transitions, and the Asian G<u>A</u>T→T, A<u>C</u>C→A, and G<u>A</u>C→T elevation was not clearly linked with sequence context features beyond a single flanking nucleotide base. However, the polymorphisms comprising profile #3 more closely matched case III, indicating that the Japanese enrichment of the *<u>A</u>C→C and T<u>A</u>T→T substitutions might be driven by a handful of 7-mer polymorphisms heterogeneous across East Asia (**Figure 3**, **Supplementary Fig. 7**).

**Figure 3:**
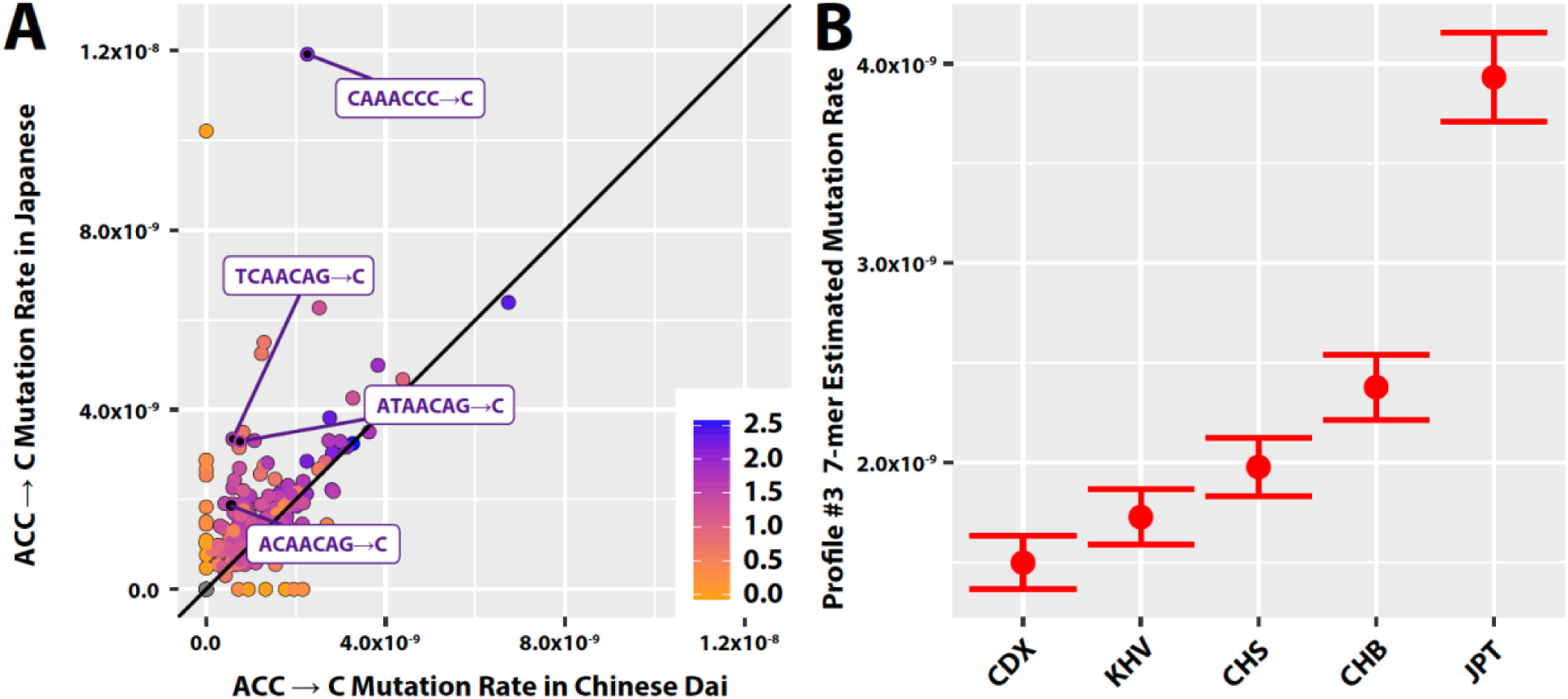
Patterns of variation within in East Asia among 7-mers from profile #3. (A) Most 7-mer expansions of A<u>A</u>C→C are the same in Chinese Dai versus Japanese, with the exception of some highly variable 7-mer polymorphism types. Polymorphisms significantly heterogeneous between Japan and Chinse Dai are labeled. **(B)** Estimated private mutation rate of the nine 7-mer polymorphism types shown in **Table 2** displayed across each East Asian subpopulation. Brackets indicate approximate 95% confidence intervals.

To explore this finding in more detail, we sought to identify the key 7-mer types underlying this 3-mer signature. To this end, we considered each of the 1,280 possible 7-mer expansions of *<u>A</u>C →C and T<u>A</u>T→T, testing for heterogeneity between Japanese individuals from Tokyo (JPT, higher profile #3 polymorphism proportion) and Chinese Dai from Xishuangbanna (CDX, lower profile #3 polymorphism proportion). We found fourteen 7-mer polymorphism types that were elevated in JPT relative to CDX (FDR-adjusted P < 0.05, **Table 2**, **Figure 3B**). Of these, ten have the shared motif XXX<u>A</u>CAG→C. Curiously, we also observed that four out of the fourteen polymorphism types were enriched on the X chromosome in East Asia, relative to the autosomes (though we note that many of the ten other substitution classes had too few observed polymorphisms on the X chromosome to justify a valid statistical test, see **Table 2**).

**Table 2:** 7-mer polymorphisms from profile #3 which are heterogenous among East Asians. Nine polymorphisms enriched in Japan compared to Chinese Dai (Chi-squared test false discovery rate (FDR) < 0.05). *Fold increase in inferred mutation rate in Japan compared to CDX. Bold p values indicate nominal significance of enrichment on the X chromosome in East Asia according to a one-sided binomial test (**Methods**). The significance values of tests for polymorphism classes with 5 or fewer observations on the X chromosome were not calculated.

| 7-mer | Fold<br>enrichment<br>in Japan* | FDR-adjusted P<br>(enrichment in<br>Japan) | Expected<br>polymorphic<br>sites on X | Observed<br>polymorphic<br>sites on X | P<br>(X enrichment) |
| --- | --- | --- | --- | --- | --- |
| TTT$\underline{A}$TTT→T | 2.14 | 6.4×10<sup>-22</sup> | 48 | 65 | **0.009** |
| CAA$\underline{A}$CCC→C | 5.28 | 1.3×10<sup>-12</sup> | 6.2 | 27 | **6.7×10<sup>-10</sup>** |
| AGT$\underline{A}$CAG→C | 16.3 | 1.3×10<sup>-7</sup> | 3.0 | 4 | - |
| TCA$\underline{A}$CAG→C | 5.64 | 9×10<sup>-4</sup> | 2.2 | 7 | **0.009** |
| ATA$\underline{A}$CAG→C | 4.37 | 0.001 | 3.3 | 7 | 0.4 |
| ATG$\underline{A}$CAG→C | 4.62 | 0.001 | 2.4 | 4 | - |
| CCC$\underline{A}$CAG→C | 2.61 | 0.001 | 4.6 | 25 | **3.3×10<sup>-11</sup>** |
| ACC$\underline{A}$CCA→C | 3.29 | 0.03 | 2.7 | 3 | - |
| AAG$\underline{A}$CAG→C | 3.20 | 0.03 | 3.1 | 4 | - |
| AAT$\underline{A}$CAG→C | 3.20 | 0.03 | 2.8 | 1 | - |
| ACA$\underline{A}$CAG→C | 4.62 | 0.03 | 2.3 | 3 | - |
| ATC$\underline{A}$CAG→C | 2.80 | 0.03 | 3.1 | 4 | - |
| GTG$\underline{A}$CAG→C | 5.86 | 0.03 | 1.2 | 1 | - |
| TTT$\underline{A}$TTA→T | 1.63 | 0.04 | 13.1 | 15 | 0.3 |

### Variable polymorphism types among broader sequence context models

Motivated by this result and previous work^3^, we next hypothesized that additional signals of mutation rate variation might be observable only in specific pentanucloetide (*i.e.*, ‘5-mer’) or 7-mer polymorphism types. If this were true, considering a broader span of sequence context would highlight novel signals of mutation rate variation not evident from 3-mer level analyses. To this end, we applied the homogeneity-testing framework described above to each of the 1,536 possible 5-mers and the 24,576 possible 7-mer polymorphism types.

Within a 5-mer sequence context, we found that 156 out of 1,535 tested polymorphism types surpassed a Bonferroni-adjusted significance threshold (**Methods and Supplementary Table 1B**; one 5-mer polymorphism was not observed a sufficient number of times in each population to justify a valid statistical test). Of these, 48 represent expansions of Europe-elevated 3-mer polymorphisms which have been highlighted by previous analyses (e.g., CT<u>C</u>CA→T, an expansion of the T<u>C</u>C→T 3-mer)^4,5.^ An additional 59 represent expansions of other 3-mers noted in **Table 1**. However, the remaining 49 significantly variable 5-mer polymorphisms involve 3-mer substitution classes that have not yet been highlighted.

We next moved to our broadest, 7-mer sequence context model. Out of 20,668 possible 7-mer substitution types with sufficient data available for a statistical test, 141 surpassed Bonferroni multiple test correction (**Table 3**, **Methods, and Supplementary Table 1C**). Of these, 119 represent expansions of polymorphisms identified at the 3-mer level, while 22 have not been previously noted. The most prominent effect at the 7-mer level was the CAA<u>A</u>CCC→C substitution (homogeneity test P_ordered_ = 3 x 10^-39^, **Table 3**) corresponding to one of the Japanese-enriched 7-mers we identified above (**Table 2**, **Figure 3A**). In total, 4 of the 22 previously unreported 7-mer polymorphisms from our significant set were among the fourteen Japanese-enriched substitution types from our earlier analysis (**Table 2**). Interestingly, the third most significant unreported 7-mer polymorphism, AAA<u>C</u>AAA→A (P_ordered_ = 1 x 10^-18^) has a similar profile within East Asia as the other profile #3 polymorphisms (**Supplementary Fig. 8**), but does not have the same 3-mer subcontext. In fact, we find that multiple 7-mer types with 3-mer subcontexts outside of *<u>A</u>C→C or T<u>A</u>T→T are enriched in Han Chinese and Japanese groups. This suggests that there exist additional 7-mer mutations comprising this signature that have not yet been discovered.

**Table 3:** Most significantly heterogenous 7-mer polymorphism types. Five most significantly heterogeneous polymorphism types at a 7-mer level, removing expansions of known 3-mer signals. Each P-value was calculated from a chi-squared test for heterogeneity across non-admixed continental groups. The Bonferroni significance for this test would be 2.5 x 10^-6^. Boldface numbers indicate a significant difference in polymorphism proportion compared with Africa (P < 1 x 10^-7^) in a pairwise chi-squared test with P_ordered_ correction. *Approximate mutation rates (per generation per site), inferred assuming a total mutation rate of 1.2 x 10^-8^ mutations per base pair per generation.

| 7-mer | Africa<br>rate* | Europe<br>rate* | South Asia<br>rate* | East Asia<br>rate* | P |
| --- | --- | --- | --- | --- | --- |
| CAA<u>A</u>CCC→C | 1 | 0.94 | 0.34 | **3.62** | $3 \times 10^{-39}$ |
| TTT<u>A</u>TTT→T | 1 | 0.84 | **0.61** | 1.04 | $2 \times 10^{-25}$ |
| TTT<u>A</u>AAA→T | 1 | 0.89 | **0.84** | **0.88** | $1 \times 10^{-21}$ |
| ATT<u>A</u>AAA→T | 1 | 0.71 | **0.76** | **0.82** | $2 \times 10^{-21}$ |
| AAA<u>C</u>AAA→A | 1 | 0.78 | **0.66** | 0.88 | $2 \times 10^{-21}$ |

Finally, two of the 7-mer polymorphism types with variable rates across populations are TTT<u>A</u>AAA→T and ATT<u>A</u>AAA→T (P_ordered_ < 2 x 10^-21^), both of which were enriched in Africa (**Figure 4**). These correspond to the 3-mer T<u>A</u>A→T, which is the 16^th^ most significant polymorphism from our 3-mer-level heterogeneity analysis (P_ordered_ = 6.2 x 10^-36^). Examining the rates of the 7-mer expansions of T<u>A</u>A→T, we find that TTT<u>A</u>AAA→T and ATT<u>A</u>AAA→T are indeed outliers among other 7-mer expansions both in terms of their African enrichment and the overall number of mutations of those types (**Figure 4B**). These results suggest that the heterogeneity we observe in proportions of TA<u>A</u>→T polymorphisms is in fact driven by an elevation of these two highly variable 7-mers in Africa.

**Figure 4:**
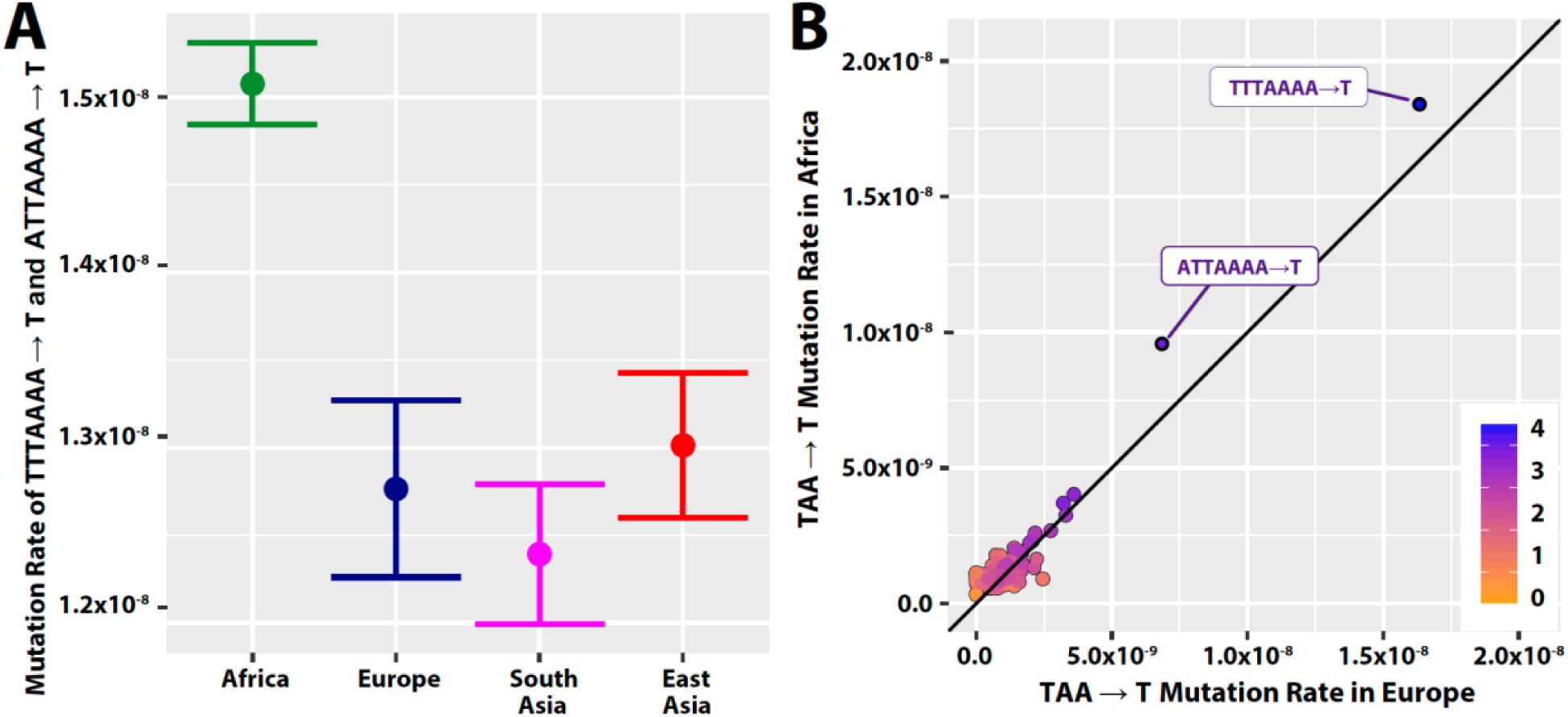
Two significantly variable 7-mer expansions of T<u>A</u>A→T. (A) Estimated pooled mutation rate of TTT<u>A</u>AAA→T and ATT<u>A</u>AAA→T across continental groups, with approximate 95% confidence intervals shown. **(B)** TTT<u>A</u>AAA→T and ATT<u>A</u>AAA→T appear to be both more variable between continental groups and more common than other 7-mer expansions of T<u>A</u>A→T. These two substitution types are the only ones from the 256 T<u>A</u>A→T expansions that are significantly different (FDR < 0.05) between Africa and Europe.

### Distribution of derived allele frequencies within mutational profiles

To better understand the mutational processes driving patterns of enrichment and depletion in certain substitution classes (**Figure 1**), we examined the distribution of derived allele frequencies (DAFs) of the polymorphisms in each profile. As in Harris and Pritchard (2017)^6^, we visualized the enrichment of each polymorphism class across DAF bins as its proportion in each DAF bin relative to its proportion across all DAF bins (**Methods**). We highlight the DAF spectra of 3 mutational profiles in **Figure 5**. The DAF spectra for all profiles in all continental groups are reported in the supplement (**Supplementary Fig. 9-13**). For profile #1, we observe the previously reported “pulse” of T<u>C</u>C→T mutations in Europe at ~1% DAF (**Figure 5A**), and see the same pulse to a lesser degree in the additional profile #1 polymorphisms identified in this work. However, we did not observe this pulse for T<u>C</u>C→T polymorphisms among private variants in South Asia (**Figure 5B**), even though this continental group does have an overall enrichment of this mutational profile. Only G<u>C</u>C→T, which in Europe has the least signature of the pulse, appears to be enriched around 1% DAF within South Asian private mutations. This suggests that though these polymorphism groups appear enriched in both Europeans and South Asians, the processes leading to the enrichment may not be the same in both continental groups. Profile #4, CpG transitions, was elevated in East and South Asian private polymorphisms. Across DAF bins, however, we observe the same pattern for all continental groups (e.g. EAS, **Figure 5C**): a relatively uniform distribution for each polymorphism class across rare variant bins, and an enrichment of all four classes in the most common DAF bin.

**Figure 5:**
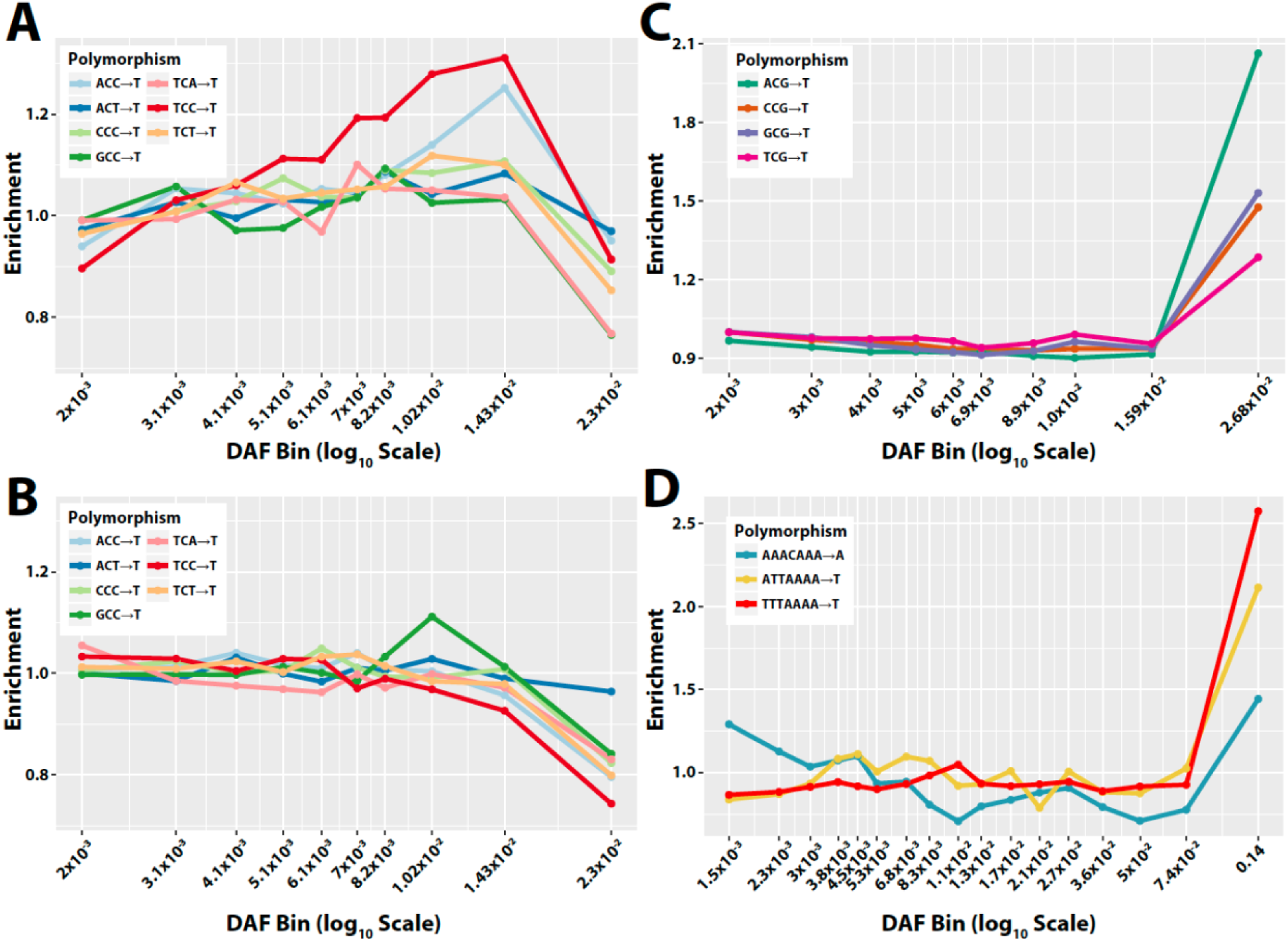
Enrichment of polymorphisms across DAF bins. The enrichment of polymorphisms in a profile across DAF bins, calculated as the proportion of mutations in a bin with a given polymorphism class, divided by the proportion of all polymorphism with that class. **(A)** Profile #1 in Europeans. **(B)** Profile #1 in South Asians. **(C)** Profile #4 in East Asians. **(D)** 7-mer contexts enriched in Africans.

Thus, it does not appear that a pulse or temporary elevation of mutation rates can explain the enrichment of CpG transitions in Asian populations. Finally, of the three 7-mer polymorphism types elevated in Africa relative to other populations (**Figure 4**), we observe an enrichment of AAA<u>C</u>AAA→A at the most rare DAF bin, which is not seen for the other two polymorphism types (**Figure 5D**). For all three 7-mers, we also observed an enrichment in the most common DAF bin, though not as strong for AAA<u>C</u>AAA→A as the other two substitution types. These results support the proposition that though the abundance of these polymorphisms is elevated in Africa, the timing of mutations leading to this enrichment may not be the same for all three classes. This suggests that more than one signature of polymorphism variation may be at work at the 7-mer sequence context level.

## DISCUSSION

In this report, we describe a number of patterns of variability in polymorphism ratios between human populations at a global scale. Whether these patterns reflect a true difference in underlying mutational processes, and what those underlying causes might be, remains unclear. Even the most prominent signature, European <u>C</u>→T, is still poorly understood: Although this group of polymorphisms appears to correlate with mutational signatures linked to ultraviolet radiation or alkylating agents in one cancer study^5,10^, evidence supporting either of these causal mechanisms is limited^5^. Moreover, while patterns of substitution rates across derived allele frequency spectra helped identify some groups of mutational changes which may have occurred in ‘pulses’ in recent history, multiple polymorphism groups lacked an obvious pulse. This collection of observations may suggests further heterogeneity in the timing of specific events that may have contributed to these changes.

Although European <u>C</u>→T enrichment is by far the most prominent signature of variation^5,6^, the large number of variable polymorphism types and the variety of patterns they follow at a global scale suggest that several different processes are at work in shaping the ratios of polymorphisms we observe. If this is correct, further scrutiny of these differences may provide an opportunity to better understand the processes that shape genomic stability and genetic change. Moreover, quantifying and modeling polymorphism patterns as accurately as possible can help us fine-tune our predictions and interpretations of single nucleotide genetic variation, potentially advancing our understanding of evolution or genetic disease.

One approach that may aid these efforts is the consideration of local genetic sequence context. Previous studies have identified different patterns of heterogeneity in polymorphism levels that can be observed between substitution classes just within different 3-mer motifs^6^, illustrating the importance of a single flanking base pair of context in shaping substitution probability. In this report, we consider a broader window of local sequence information, noting that while certain signatures are fully explained by a single flanking base pair or context, others appear to vary with sequence context up to 2-3 base pairs from the locus of substitution (**Figures 2, 3B, and 4C**). We report that ten of the fourteen heterogeneous 7-mers between Chinese Dai and Japanese in profile #3 contain the 7-mer motif XXX<u>A</u>CAG→C (**Table 2**). In addition, we find that the apparent elevation of T<u>A</u>A→T 3-mer polymorphisms between Africa and Europe may in fact be driven by a strong enrichment of substitutions within WTT<u>A</u>AAA contexts (where ‘W’ represents a weak ‘A’ or ‘T’ base), which also appear to segregate more substitutions than other T<u>A</u>A contexts (**Figures 4B and 4C**).

There are some limitations to note in our study. First, is sample size: while broader sequence context models can capture more information, they can also require much more total genetic data to be sufficiently well-powered for our statistical analysis. This is made especially difficult because asking comprehensive questions about global mutation rate patterns requires a large and ethnically diverse dataset of genetic variation, which are only recently becoming available. Additional, deeply sequenced samples from diverse populations would be ideal for further targeted hypothesis testing, validation, and improvement of the mathematical models designed to capture this variability. For example, in this report, we noted evidence suggesting that East Asian heterogeneity in *<u>A</u>C→C and T<u>A</u>T→T mutations may be strongest on the X chromosome (**Table 2**). Given this observation, it may be informative to examine the dispersion of these polymorphisms across the X-chromosome, since a genetic variant responsible for an increase in mutation rate is likely to be found in a genetic context with high rates of polymorphism^11^. Unfortunately, however, a problem of power quickly emerges, since the total number of polymorphisms of any 7-mer context observed on the X chromosome is still relatively small. As a result, analyses regarding this signature may be difficult until a larger amount of East Asian genetic data is made available.

A second complication is that signals of polymorphism enrichment from population-level data may reflect some contemporary and some ancestral mutation rate variation. As such, it is not immediately clear whether the biological mechanisms driving these phenomena are still active today. Measurements of enrichment of these polymorphism types in ancient DNA and across allele frequency bins suggest that the previously reported European signal (profile #1) may correspond to an ancestral increase in the rate of certain <u>C</u>→T mutations ~15,000 years ago, which may have subsided ~2,000 years ago^5,6^. In this study, we examined the distribution of polymorphism enrichment across allele frequencies, allowing us to hypothesize about the timing of these events across continental groups (**Figure 5**). Further analyses might consider the timing of polymorphism enrichment using direct estimates of allele age, (rather than allele frequency as a proxy for age). This may help us better piece together the timescale over which mutation rates may have changed^6^.

It is likely that further investigation will reveal details of mechanism, evolutionary timing, and genome-wide or subpopulation-level patterns in polymorphism variation, and our report here is by no means exhaustive. We detail evidence suggesting that polymorphism variation acts in a variety of ways across human populations based on local sequence context cues at varying distances from the mutated locus. While some of these signals manifest at the 3-mer level, consideration of a broader context brings new patterns of variation to light.

## METHODS

### Compilation of private variant sets

Variants from the 1,000 Genomes Project release (downloaded 02/26/2016, phase 3)^8^ were filtered to include single nucleotide polymorphisms (SNPs), excluding multiallelic variants, in/dels, and any variants with a filter tag other than “PASS”. Based on the exclusion criteria from previous work, we also omitted variants in coding regions, centromeres, telomeres and additional sections of the genome predicted to have low accessibility^3^. From this set, we considered variants with a minor allele count 2 or greater across all samples. Although including singleton variants (those observed only once in the dataset) in theory would provide more information about the recent *de novo* mutation rate, previous efforts to analyze human polymorphism variation with singletons have proven difficult to replicate^4^, so we opted to exclude them from our analyses.

From variants that passed our filters, we compiled lists of variants ‘private’ to each non-admixed continental group from the dataset: Africans from Africa (AFR), Europeans (EUR), East Asians (EAS), and South Asians (SAS). We defined a SNP as ‘private’ to a continental group if it is observed in that group, but not in each of the other three. For all analyses, Americans of African Ancestry in Southwest USA (ASW) and African Carribeans from Barbados (ACB) were considered to be admixed American populations rather than ancestral African groups.

For subpopulation-level analyses, we passed along private polymorphisms defined for each continental group into each respective subpopulation list. For example, a polymorphism which was private to AFR and observed in both Kenya and Gambia would be added to the subpopulation lists for both LWK (Luhya in Webuye, Kenya) and GWD (Gambians in Western Divisions in Gambia).

American admixed variant lists were compiled from all SNPs which were present in an American subpopulation but not present in more than one ancestral continental group. All filtration steps were carried out using vcftools and the vcf-isec tool (v0.1.12b)^12^.

From each continental or subpopulation list, we tallied counts of private variants by increasingly broader windows of sequence context, including one, two, and three flanking nucleotides (*i.e.*, 3-mer, 5-mer, and 7-mer context, respectively). During this process, mutation classes were ‘folded’ to include its reverse-complimentary equivalent (*e.g.*, T<u>C</u>C→T and G<u>G</u>A→A were considered together). Code utilized for each step is available online (github.com/raikens1/mutation_rate).

### Statistical comparison with homogeneity test

To replicate previous work by Harris and Pritchard^4,6^, we first performed pairwise chi-squared comparisons of polymorphism count between each possible pair of populations for each 3-mer polymorphism type (**Supplementary Fig. 1**). Next, to partially relieve the multiple testing burden of six pairwise population comparisons over each possible mutation type, we performed a single test using a 2x4 cell contingency table which included polymorphism counts for each continental groups. The resulting chi-squared test for homogeneity from this table has three degrees of freedom. An issue with performing this type of analysis across all substitution classes is that the P-values from these tests are non-independent: a polymorphism that is profoundly heterogeneous across populations may alter the proportions of other polymorphism types. For these reasons, we used the *P_ordered_* correction as previously described^6^. Using this procedure, each polymorphism type is initially tested and ranked according to increasing significance based on a simple homogeneity test using all the data. A corrected p-value is then calculated for each polymorphism. To do this, the least significant polymorphism is assigned its original p-value using all of the data. After this, the p-value for the *i*^th^ least significant polymorphism type is recalculated using a homogeneity test with only the data for the *i* least significantly variable polymorphisms from the initial ranking. All chi-squared comparisons were performed using the chisq.test function in R (v3.4.0), and significance thresholds were determined based on a Bonferroni correction with a nominal error rate (α = 0.05).

### Mutation rate inference

The probability of observing a given polymorphism in a population is determined by a composite of mutation rate, demography, sample size, and other factors^13^. To facilitate comparisons across populations, we calculated a mutation rate, calibrated to the average *de novo* mutation rate estimated by Kong et al^9^. Assuming all populations have a total mutation rate of 1.2 x 10^-8^, we inferred the mutation rate of a specific type (say T<u>C</u>C→T) as

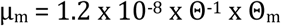

Where μ_m_ represents the inferred private germline T<u>C</u>C→T mutation rate per generation per site, Θ_m_ represents the proportion of all T<u>C</u>C sites in the genome with private C/T polymorphism in the population, and Θ represents the total proportion of all sites of any type in the genome which are private polymorphisms in the population. It can be shown that this formulation of μ_m_ gives an overall genome wide mutation rate of 1.2 x 10^-8^ when all mutation types are pooled. 95% confidence intervals for μ_m_ were calculated using the normal approximation to the binomial, assuming the variance in our estimate of Θ^−1^ to be approximately zero.

### Clustering polymorphism types

We used the heatmaps 2 hierarchical clustering methods from the basic stats package in R (v3.4.0) in order to heuristically identify mutation types that vary in similar ways across the globe. In doing so, we defined the “profile” of a mutation *m* across a set of populations as a vector of the inferred mutation rate of *m* in each population. Each pattern of rates across populations was normalized by fold difference above or below the mean rate for that profile. We used Euclidean distance to construct each heatmap, selected because these methods gave the most clearly interpretable results and agreed the most closely with previous work^4–6^

### Testing for enrichment of profile #3 on the X chromosome

In order to test the highly variable polymorphism types for enrichment on the X chromosome (**Table 2**), we used a one-sided binomial test to determine whether the observed proportion of privately polymorphic sites on the X chromosome was greater than *p_0_*, the expected proportion under the null hypothesis. For demographic and sampling reasons, we expect to observe fewer polymorphic sites of any given type on the X chromosome than on the autosomes, even if the mutation rate of that polymorphism type is identical across chromosomes. Thus, to estimate *p_0_*, we first needed to calculate the ratio, *ξ* of X-chromosome to autosomal substitution probability across all other 7-mer types. We then used this as a scaling factor, estimating *p_0_* as *ξp_A_*, where *P_A_* represents the maximum likelihood estimate for the substitution probability for that polymorphism across all autosomes.

### Enrichment across DAF bins

To identify the derived allele frequency of mutations private to each continental group, we used the ancestral allele calls reported in the 1000 Genomes Phase 3 release VCF files. These ancestral calls are originally from the Ensembl Compara release 59^14^. For the purposes of this specific analysis, we omitted variants that did not have a high-confidence ancestral state call, and variants on the X chromosome. For each continental group, we binned variants by identifying the quantiles of DAFs by 5% increments, and collapsing to only quantiles with unique values. In this way we tried to ensure that each bin had a reasonable amount of data, though not all bins were the same size. We calculated enrichment of polymorphisms within a sequence context in each bin as the proportion of polymorphisms in that bin with that context, divided by the proportion of all polymorphisms across all bins with the given context.

## ACKNOWLEDGEMENTS

R.C.A. is thankful for the guidance and feedback of Julian Segert, Elizabeth Vallen, Nick Kaplinsky, Bradley Davidson, and Ameet Soni. The authors would also like to thank Varun Aggarwala and Onur Yörük, who offered helpful advice on the implementation of 5-mer and 7-mer sequence context analyses. B.F.V. is grateful for support from the US National Institutes of Health/National Institute of Diabetes and Digestive and Kidney Disorders (R01DK101478). KEJ was supported in part by NIH training grant T32GM008216.

## AUTHOR CONTRIBUTION

R.C.A., K.E.J., and B.F.V. conceived, designed, and performed experiments developed methods, analyzed data, and wrote the manuscript. B.F.V. supervised the research.

